# Increasing Pyrethroids and DDT Resistance and kdr Mutation in *Anopheles gambiae s.l.* from Sokoto, North-West Nigeria

**DOI:** 10.64898/2026.04.27.720970

**Authors:** Usman Salisu Batagarawa, Musa Abubakar Yalwa, Abubakar Sani, Muhammad Awwal Abdullahi, Amir Yakubu Gobir, Yusra Ahmad Adamu, Bilkisu Kabir Rawayau, Sulaiman Lawal Dara, Victoria Olawole Temitope, Vanessa Pius Godiya, Ayman Abdullahi Muhammad, Abdulmalik Sani, Jamilu Ibrahim, Andrew Onu, Iliya S. Ndams, Abdullahi B. Sallau, Mohammed Nasir Shuaibu, Jun Hang, Yahaya Muhammad Abdullahi

## Abstract

**Background:** *Anopheles gambiae sensu lato* (*s.l*.) is the primary vector of malaria in sub-Saharan Africa. Although insecticide-based vector control has been central to prevention, the widespread emergence of insecticide resistance poses a serious biological threat to control efforts. Effective resistance monitoring is essential for sustaining vector control but remains highly limited in malaria-endemic hotspots. Here, we assessed pyrethroid and DDT resistance intensity and the frequency of the L1014F knockdown resistance (kdr) mutation in *Anopheles gambiae* s.l. populations from Sokoto, north-western Nigeria.

**Methodology:** Resistance status and intensity to five insecticides were determined in adult *Anopheles* reared from larvae collected in 2021 and 2022 using the World Health Organization (WHO) tube test and Centers for Disease Control and Prevention (CDC) bottle bioassay, respectively. A subset of resistant mosquitoes was analyzed using PCR-based diagnostic assays to identify species within the *Anopheles gambiae* complex and to genotype for the West African *kdr* mutation (L1014F).

**Results:** High knockdown times (KDT) were observed, with KDT50 ranging from 38 to 91 minutes and KDT95 from 104 to 678 minutes, indicating increased resistance levels across all insecticides. In 2021, resistance was detected to DDT, lambda-cyhalothrin, and permethrin, while susceptibility to alpha-cypermethrin (98%) and suspected resistance to deltamethrin (91%) were recorded. In 2022, a general increase in resistance to all insecticides was observed, with mortality rates ranging from 41% to 81%. High resistance intensity was observed against DDT, while permethrin and alpha-cypermethrin exhibited low resistance intensity in both years, failing to reach 10x the diagnostic dose. Deltamethrin and lambda-cyhalothrin showed low to moderate resistance intensity. The 1014F *kdr* mutant genotype was widely distributed (68.1%) across species and years. Allele frequencies were higher in *An. gambiae s.s*. (0.83) than in *An. arabiensis* (0.71), with significant deviations from Hardy-Weinberg equilibrium (p < 0.05), except for *An. gambiae s.s*. in 2021 (p = 0.7).

**Conclusion:** These findings reveal a concerning increase in key insecticide resistance among *Anopheles* populations in Sokoto, underpinned by strong genetic mechanisms. This underscores the urgent need for integrated vector management strategies to sustain effective vector control efforts in the region.

## Background

Malaria and lymphatic filariasis remain major public health burdens worldwide, transmitted predominantly by *Anopheles* mosquitoes, which serve as the principal vectors of *Plasmodium* parasites and *Wuchereria bancrofti*, respectively (1,2). In Nigeria, about 30 *Anopheles* species are present (3); among these, members of the *Anopheles gambiae* complex (*An. gambiae s*.*s*., *An. arabiensis*, and *An. coluzzii*) are the primary malaria vectors, while *An. funestus* acts as a secondary vector (4). Malaria continues to be a major global health concern, with an estimated 263 million cases and 597,000 deaths recorded in 2023. It is estimated that over half of the world’s population live at risk of malaria or other febrile vector-borne illnesses (5). Nigeria bears the heaviest burden globally, accounting for 26% of all malaria cases, equivalent to the combined burden of the next four highest-burden countries, namely the Democratic Republic of the Congo, Uganda, Ethiopia, and Mozambique. Nigeria also recorded the highest malaria mortality (30.9%) and accounted for 39.3% of global malaria deaths among children under five (5). The northwest region of the country has the highest prevalence at 33.8% (6), with Sokoto State (36%) ranking second nationally, after neighboring Kebbi State (49%) (7). In all, the entire Nigerian population remains at risk of malaria infection, with approximately 76% living in high-transmission areas and the remainder in low-transmission zones (8). In response, several national initiatives have been launched in line with global malaria eradication goals, including the National Malaria Strategic Plan (NMSP), the Nigeria End Malaria Council (NEMC), and the National Malaria Elimination Program (NMEP), all targeting eradication by 2030 (9). Central to these strategies is vector control, which remains the cornerstone of malaria prevention.

Vector control interventions are primarily implemented through a combination of long-lasting insecticidal nets (LLINs) and indoor residual spraying (IRS), both of which rely heavily on chemical insecticides (10,11). Among these, pyrethroids—predominantly permethrin and deltamethrin—are the most widely used, accounting for 89.9% of insecticides applied for standard spray coverage in Africa (12). Their popularity stems from their strong residual effects, low mammalian toxicity, rapid knockdown action, and high toxicity to mosquitoes. In 2023 alone, 255 million insecticide-treated nets (ITNs), the primary vector control tool, were distributed in malaria-endemic countries, including 24 million in Nigeria (5). Although these large-scale interventions have contributed significantly to malaria control, they have also exerted strong selection pressure on mosquito populations, driving the emergence of insecticide resistance. This resistance poses a major biological threat to malaria control, enabling mosquitoes to survive exposure to all major insecticide classes, particularly pyrethroids (5,13).

The emergence and spread of insecticide resistance are primarily driven by two mechanisms: target-site insensitivity (mutation) and metabolic resistance. The most widespread and well-characterized form is target-site resistance, commonly referred to as knockdown resistance (kdr) (14,15). This mechanism results from genetic mutations in the voltage-gated sodium channel (VGSC) gene, the primary target of pyrethroid and Dichlorodiphenyltrichloroethane (DDT) insecticides. Specific point mutations in the VGSC confer varying levels of resistance by reducing the channel’s sensitivity to insecticides. The most notable are the L1014F (kdr-west) and L1014S (kdr-east) substitutions, where leucine (L) at amino acid residue (AA) 1014 is replaced by phenylalanine (F) or serine (S), respectively. The kdr-west (kdr-w) L1014F mutation is geographically restricted and highly prevalent in West Africa, likely driven by ecological conditions and intense selection pressure from the widespread DDT and pyrethroid use (16– 18). Other substitutions at AA1014, as well as novel non-1014 VGSC mutations, are increasingly being reported, either alone or in combination with L1014 mutations, further strengthening resistance (19–22).

Insecticide resistance has long been a major challenge to malaria control in Nigeria. It was first reported in the old Sokoto region as early as the mid-1950s, before the country’s independence, largely attributed to intensive insecticide use (23,24). Today, resistance to all major pyrethroids and DDT, along with associated molecular mechanisms, is widespread across Africa and consistently reported in all six geopolitical zones of Nigeria (15,25–44) However, most of these reports originate from southern Nigeria, and clear regional variations exist. In northwestern Nigeria, the kdr-w mutation has also been documented (29–31,37,45), but information from Sokoto State remains limited. Importantly, resistance status and *kdr* allele frequencies vary not only across regions but also between locations within the same region, and resistance intensity may change over time even within a single locality.

To address these challenges, insecticide resistance monitoring in Nigeria has been strengthened in recent years, particularly through annual surveillance activities conducted by the U.S. President’s Malaria Initiative (PMI) VectorLink project across the country’s five ecological zones, with Sokoto State serving as a sentinel site (46,47). In line with World Health Organization (WHO) recommendations, regular monitoring of resistance profiles in highly endemic regions is essential for guiding the choice of vector control tools (48,49). Effective resistance management therefore requires not only continuous monitoring of resistance status and intensity but also characterization of the underlying mechanisms. As such, the present study investigates the resistance status and intensity of *Anopheles gambiae* s.l. populations from Sokoto, northwestern Nigeria, against key pyrethroid and DDT insecticides, and determines the frequency of the L1014F kdr mutation in sibling species.

## Methods

### Study Area and Design

*Anopheles* mosquitoes were cross-sectionally sampled from Usmanu Danfodiyo University Sokoto (UDUS) main campus and surrounding localities **(Fig. 1)**. Sites were selected based on proximity to human activity areas, small-scale cultivation and livestock, and the availability of breeding habitats, including open drainages, stagnant pools formed by water spillage from wells or boreholes, and rainwater pools. The university is surrounded by several rural communities where residents primarily practice farming and animal rearing, often using chemical pesticides and fertilizers. UDUS is located in Wamako Local Government Area, approximately 7 km south of Sokoto city, in Sokoto State, northwestern Nigeria. Its coordinates are approximately 13.1°N, 5.2°E, at an elevation of about 309m above sea level. The area lies within the Sudan savanna ecological zone where mean daily temperatures range between 15 °C and 40 °C, characterized by a semi-arid climate with a rainy season from June to October (annual rainfall: 500–800 mm) and a prolonged dry season from November to April/May.

**Fig. 1:**
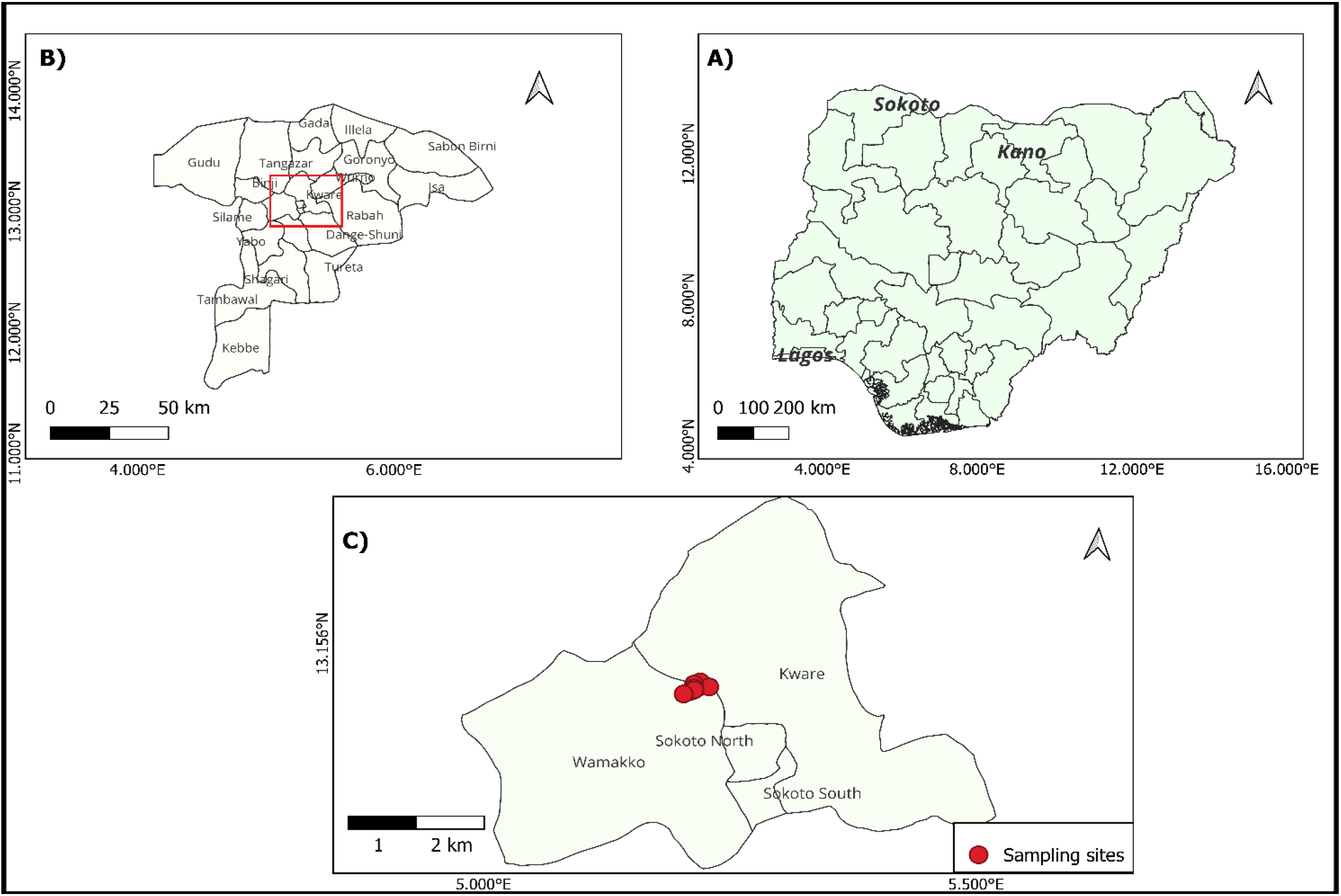
Geographic location of the study area and sampling sites in Sokoto, northwestern Nigeria. **A)** Map of Nigeria showing the location of Sokoto State. **B)** Administrative map of Sokoto State displaying local government areas, with the study area highlighted. **C)** Detailed map of the study area indicating sampling sites within major parts of Sokoto metropolis (Sokoto North, Sokoto South, Wamakko, and Kware Local Government Areas).

### Mosquito Larval Collections and Rearing

Immature stages of *Anopheles* mosquitoes (larvae and pupae) were collected from various *Anopheles* larval habitats within the study area during the rainy seasons of June–September 2021 and May–August 2022. Collections were made using the standard dipping method (50) with a 350 ml larval dipper (Bio Quip Products Inc., Gardena, California, USA), transferred into plastic containers, filtered through sieves, and transported to the insectary. Larvae were reared to adulthood under standard insectary conditions (27 ± 2 °C, 75 ± 10% relative humidity, and a 12 h:12 h light/dark photoperiod) and fed with crushed plain biscuits (51,52). Emerged adults were maintained on 10% sugar solution soaked in cotton pads (53).

### WHO Insecticide Susceptibility Bioassay

Phenotypic insecticide susceptibility was assessed following the standard WHO tube bioassay protocol (54). Briefly, blood-unfed 2–5-day-old female *An. gambiae* s.l. were exposed to WHO-prepared insecticide-impregnated papers at the following discriminating concentrations: permethrin (0.75%), deltamethrin (0.05%), alpha-cypermethrin (0.05%), lambda-cyhalothrin (0.05%), and DDT (4%). For each insecticide, about 20–25 mosquitoes were exposed in four replicate tubes, with an additional control tube containing only carrier oil. Mosquitoes were exposed for 1 hour, and knockdown was recorded every 10 minutes. After exposure, mosquitoes were transferred to clean holding tubes with untreated papers and maintained on 10% sugar solution ad libitum soaked cotton wools as a food source. Final mortality was recorded 24 hours post-exposure. The identity of dead (susceptible) and surviving (resistant) mosquitoes were confirmed morphologically using taxonomic keys from (55) and the Walter Reed Biosystematics Unit (WRBU) online pictorial identification key (https://wrbu.si.edu/vectorspecies/keys) for Afrotropical mosquitoes. Specimens were stored in labeled Eppendorf tubes with blue silica gel beads for molecular analysis.

### CDC Resistance Intensity Bioassay

Resistance intensity was assessed using the CDC bottle bioassay (56), with modifications following (42) Adult females were exposed to increasing multiples of the diagnostic concentrations (1×, 5×, 10×): permethrin (21.5 µg/bottle), deltamethrin (12.5 µg/bottle), alpha-cypermethrin (12.5 µg/bottle), lambda-cyhalothrin (12.5 µg/bottle), and DDT (100 µg/bottle). Groups of 20–25 females were exposed in 250 mL Wheaton bottles coated with the respective insecticide doses. Following exposure, mosquitoes were transferred to net-covered disposable paper cups, provided with 10% sugar solution, and ethanol-coated bottles used as controls. Mortality was recorded after 24 hours, and percentage mortality was calculated.

### DNA Extraction

A subset of resistant mosquitoes was selected for molecular analysis to identify sibling species and genotype the kdr-w L1014F mutation. Genomic DNA was extracted using the DNeasy Blood & Tissue Kit (Qiagen Cat. No. 69504; Hilden, Germany) following the manufacturer’s protocol, and stored at -20 °C until PCR analysis.

### Molecular Species Identification

Species within the *Anopheles gambiae* complex were identified using a PCR-based method targeting ribosomal DNA intergenic spacer (IGS) regions (57), with minor modifications. Multiplex PCR was performed on Applied Biosystems GeneAmp™ PCR System 9700 Thermal Cycler using a universal (UN) forward primer and three species-specific reverse primers: GA (*An. gambiae s*.*s*.), AR (*An. arabiensis*), and QD (*An. quadriannulatus*) (**Table 1**). Reactions were carried out in a total volume of 25 µl using the Qiagen TopTaq Master Mix Kit (Qiagen Cat. No. 200403; Hilden, Germany), containing 12.5 µl of 1× TopTaq Master Mix, 0.5 µl of each primer (final concentration of 0.2 µM), 2.5 µl of CoralLoad, 4.5 µl of nuclease-free water (NFW), and 5 µl of template DNA or NFW (for negative control). PCR cycling conditions: initial denaturation at 94 °C for 3 min, followed by 40 cycles of denaturation at 94 °C for 45 s, annealing at 50 °C for 30 s, and extension at 72 °C for 45 s, with a final extension of 10 min at 72 °C. PCR products were separated on 2% agarose gels stained with ethidium bromide. Species were identified by expected band sizes: 390 bp (*An. gambiae s*.*s*.), 315 bp (*An. arabiensis*), and 153 bp (*An. quadriannulatus*).

**Table 1:**
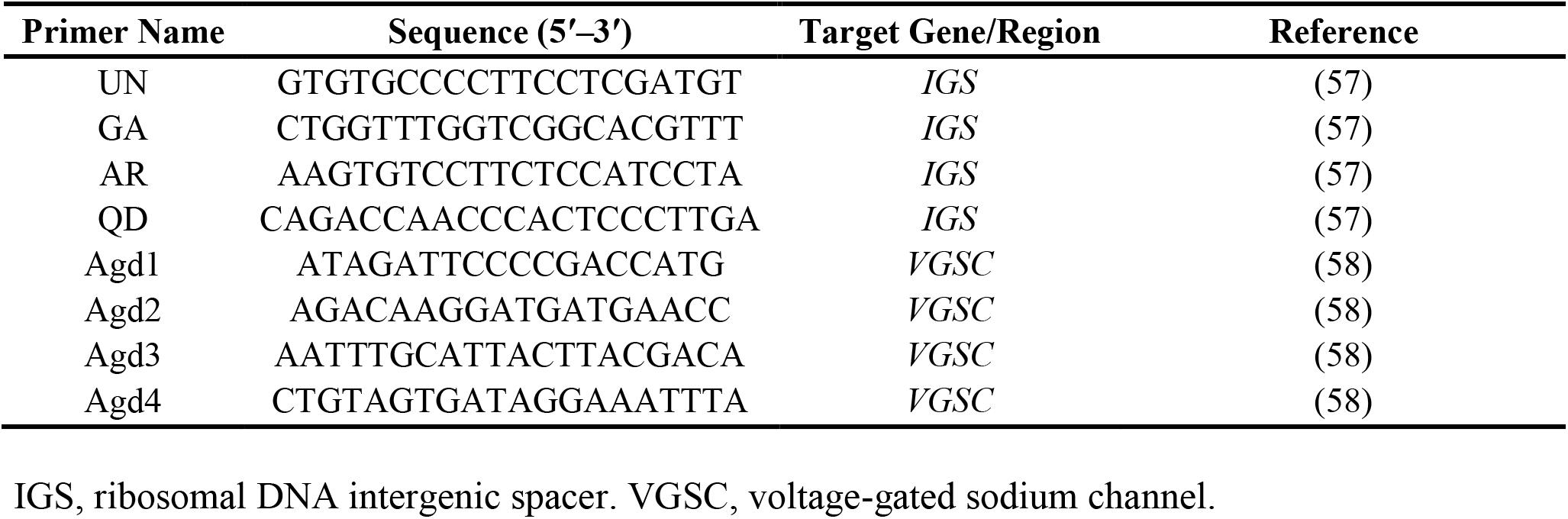
Primers used for PCR in species identification and kdr genotyping.

### Allele-Specific PCR Genotyping L1014F kdr Allele

The L1014F mutation in the VGSC gene was genotyped using allele-specific PCR (AS-PCR) following the protocol described by (58). Four primers, forward primers Agd1 and Agd4 and reverse primers Agd 2 and Agd3 (**Table 1**), were used in 25 μl PCR reactions. PCR cycling conditions were the same as above, except annealing at 54 °C for 30 s. The expected products included: 293 bp (internal control, Agd1/Agd2), 195 bp (resistant allele, Agd1/Agd3), and 137 bp (susceptible allele, Agd4/Agd2). Products were separated on 2% agarose gels stained with ethidium bromide, and genotypes were assigned based on banding patterns: homozygous susceptible (L/L), homozygous resistant (F/F), and heterozygous (L/F).

### Statistical Analysis

Bioassay results were expressed as percentage mortalities to assess resistance status and intensity. Mortality data were corrected using Abbott’s formula (Abbott, 1925) when control mortality was 5–20%, while assays with control mortality of >20% were discarded. Knockdown times (KDT50 and KDT95, the time needed to knock down 50% and 95% of mosquitoes respectively), slopes, chi-square values, and *p*-values were estimated using log-time probit analysis with LdP Line software (Ehabsoft), according to (59).

Resistance status was classified according to (54) criteria:

- Susceptible: 98–100% mortality
- Suspected resistance: 90–97% mortality
- Resistant: <90% mortality

Resistance intensity was further classified per (54) guidelines:

- Low intensity: 98–100% mortality at 5× concentration
- Moderate intensity: <98% mortality at 5× but 98–100% at 10×
- High intensity: <98% mortality at 10×

Genotype frequencies of the L1014F mutation were calculated, and allele frequencies were derived. Hardy–Weinberg equilibrium (H-WE) was tested by comparing observed and expected genotype distributions.

## Results

### Mosquito samples for adult insecticide susceptibility bioassays

A total of 7,735 adult mosquitoes successfully emerged, all of which (100%) were morphologically identified as *Anopheles gambiae* s.l. Of these, 3,012 were collected in 2021 and 4,723 in 2022 from shallow pools, puddles, and blocked drainages. From the emerged mosquitoes, 822 *An. gambiae* s.l., consisting of 422 from 2021 and 400 from 2022, were exposed to four pyrethroids and DDT using the standard WHO tube bioassay.

### Decline in knockdown efficiency of insecticides

A decline in knockdown response across all insecticides tested was observed between 2021 and 2022 and is further analyzed below. The knockdown profiles within and after the 1-hour exposure period are presented in **Fig. 2 and Fig. 3**, respectively. Knockdown rates increased progressively with exposure time; however, marked differences were evident among insecticides and between years, particularly beyond 40 minutes of exposure. Overall, alpha-cypermethrin consistently exhibited the strongest knockdown effect, producing 88.8% and 72.5% knockdown after 60 minutes of exposure in 2021 and 2022, respectively. In contrast, DDT produced the slowest knockdown response, with approximately 40% knockdown in both years, consistent with widespread resistance.

**Fig. 2:**
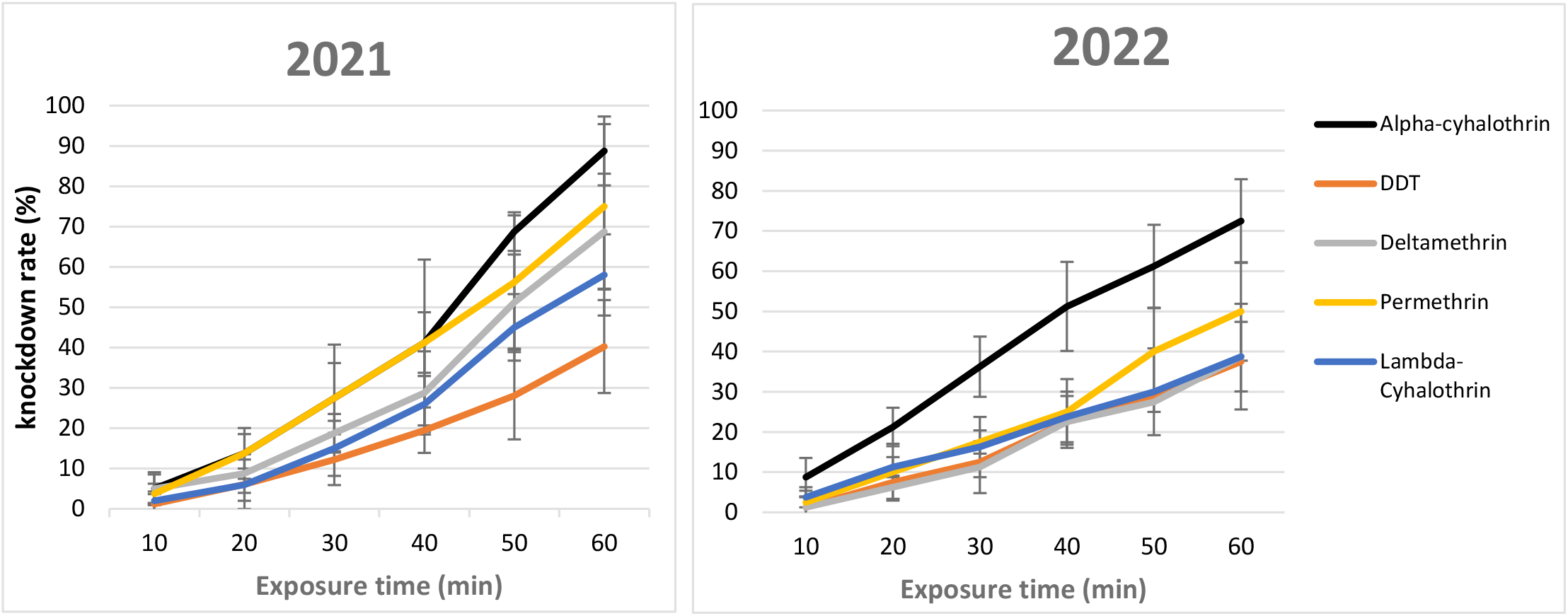
Evolution of *An. gambiae* s.l. knockdown rates during the 1-hour exposure period to pyrethroid and DDT insecticides. The chart illustrates inter-annual variation between 2021 and 2022, with each line representing the mean knockdown (%) response to a specific insecticide measured at 10-minute intervals. For each insecticide, an average of 20–25 mosquitoes were exposed in four replicate tubes. Error bars indicate standard deviations of replicate bioassays.

**Fig. 3:**
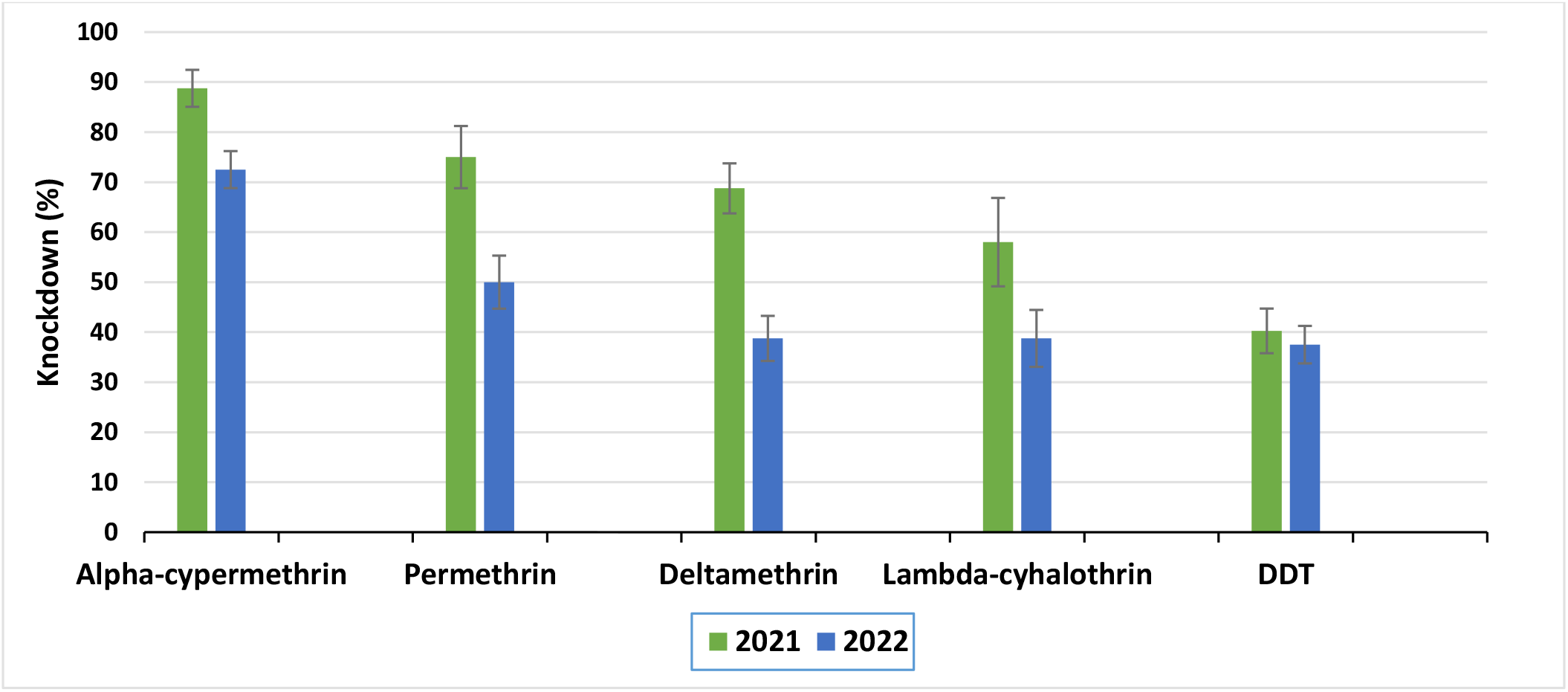
Mean knockdown percentage of *An. gambiae* s.l. after 1-hour insecticide exposure. Bars represent the inter-annual mean knockdown response at the end of a 60-minute exposure for each insecticide, arranged in knockdown (%) order: Alpha-cypermethrin, Permethrin, Deltamethrin, Lambda-cyhalothrin, and DDT. Error bars indicate the standard error of four replicate tubes in WHO tube bioassays.

The most substantial and significant (p < 0.05) reductions were observed for deltamethrin (up to 4-fold lower at early time points and 1.8-fold at 60 min), permethrin, and lambda-cyhalothrin (both showing a 1.5-fold decrease at 60 min). A moderate reduction was observed for alpha-cypermethrin (1.2-fold at 60 min), while DDT knockdown showed minimal variation, remaining uniformly low and largely unchanged. Furthermore, the fold reduction in knockdown response between 2021 and 2022 across exposure times (**Fig. 4**) illustrates higher knockdown rates at all time points for deltamethrin and permethrin in 2021. A decline was observed only at 50 minutes for lambda-cyhalothrin and alpha cypermethrin, whereas DDT exhibited only a minimal decrease.

**Fig. 4:**
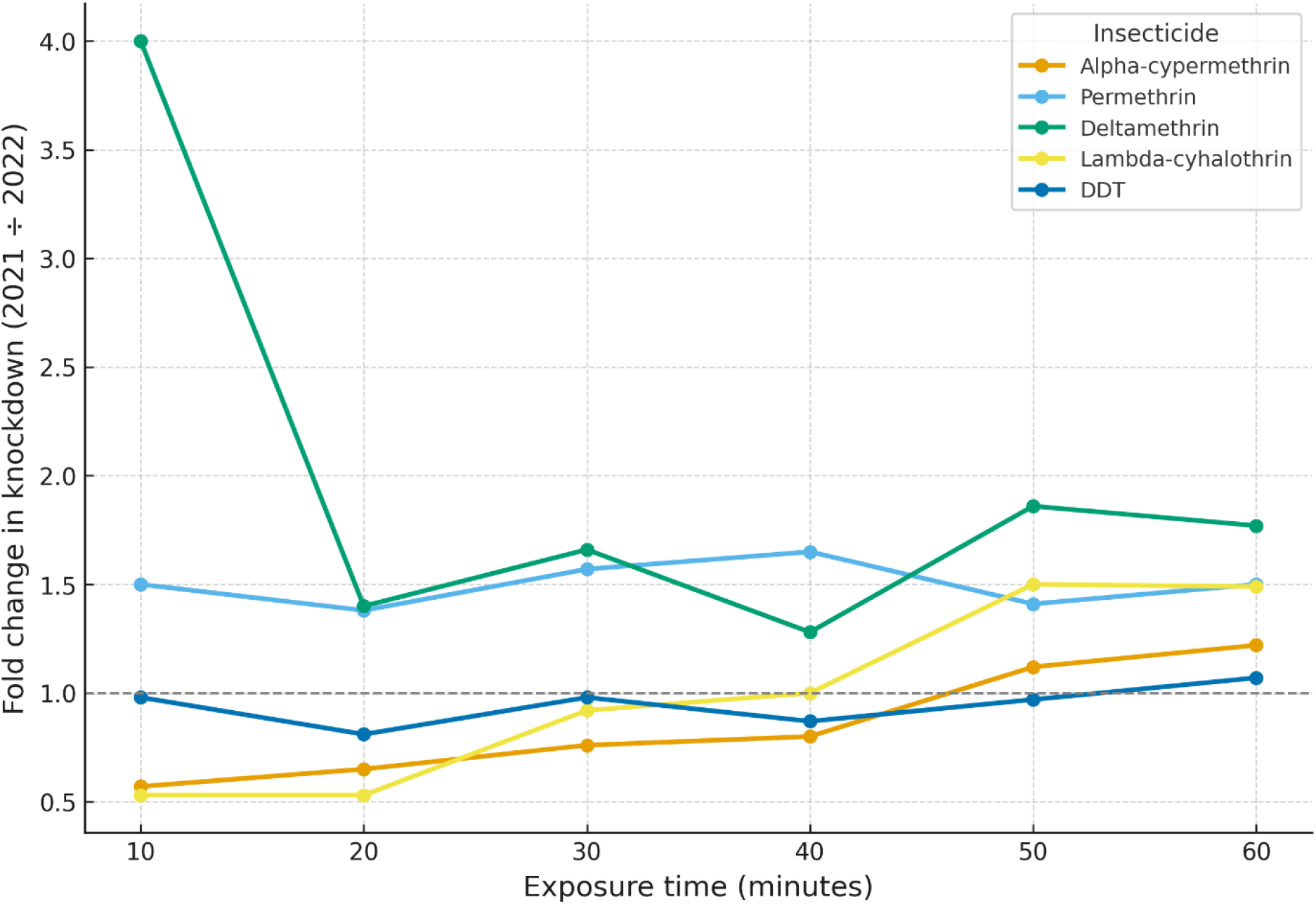
Fold reduction in knockdown response of An. gambiae s.l. between 2021 and 2022 across different exposure times for five insecticides. The dashed line at 1.0 represents “no change” and serves as the reference line, while each line corresponds to a specific insecticide. Values above the reference line indicate a decrease in knockdown in 2022 relative to 2021, whereas values below it indicates higher knockdown in 2022.

### Knockdown Time Comparisons

Results from log-time probit analysis of *Anopheles* populations exposed to pyrethroids and DDT are summarized in **Table 2**. A substantial variation and increase were observed across all insecticides and between years. In 2021, KDT50 values for the four pyrethroids ranged from 38.15 to 56.35 min, while in 2022 they ranged from 38.36 to 91.69 min. The corresponding KDT95 values ranged from 104.01 to 173.88 min in 2021, and from 167.56 to 678.04 min in 2022. Alpha-cypermethrin consistently produced the lowest knockdown times in both years. Although its KDT50 remained relatively stable, its KDT95 increased 1.6-fold 2022. In contrast, lambda-cyhalothrin showed the slowest knockdown effects, recording the highest KDT50 and KDT95 values in both years, exceeding even those of DDT in 2022. However, the KDT95 values of lambda-cyhalothrin and deltamethrin were comparable in 2021.

**Table 2:** Knockdown times (KDT50 and KDT95) of *An. gambiae* s.l. during 1-hour exposure to Pyrethroids and DDT.

| Year | Insecticides | No. exposed | KDT50 (95% CI) | KDT95 (95% CI) | $\chi^2$ (p) | Slope |
| --- | --- | --- | --- | --- | --- | --- |
| 2021 | Alpha-cypermethrin | 80 | 38.15 (32.36-45.55) | 104.01 (76.07-197.89) | 5.02 (0.29) | 3.78± 0.71 |
|  | DDT | 82 | 80.14 (66.06-111.35) | 338.23 (204.71-848.42) | 0.82 (0.94) | 2.63± 0.41 |
|  | Deltamethrin | 80 | 50.62 (41.73-84.70) | 173.86 (179.63-933.56) | 14.18 (0.01) | 3.07±0.35 |
|  | Lambda-cyhalothrin | 100 | 56.35 (47.11-78.44) | 173.88 (109.68-532.63) | 6.19 (0.19) | 3.36±0.37 |
|  | Permethrin | 80 | 43.13 (39.34-47.93) | 139.64 (109.77-199.44) | 4.26 (0.37) | 3.23± 0.34 |
| 2022 | Alpha-cypermethrin | 80 | 38.36 (34.40-43.29) | 167.56 (124.49-262.25) | 1.48 (0.83) | 2.57±0.28 |
|  | DDT | 80 | 83.17 (67.27-120.35) | 395.62 (226.91-1117.86) | 0.33 (0.99) | 2.43±0.38 |
|  | Deltamethrin | 80 | 81.25 (66.48-114.83) | 354.09 (210.20-931.35) | 0.71 (0.95) | 2.57±0.40 |
|  | Lambda-cyhalothrin | 80 | 91.69 (70.07-150.33) | 678.04 (323.89-2905.20) | 0.5 (0.97) | 1.89±0.32 |
|  | Permethrin | 80 | 64.53 (55.39-81.36) | 277.23 (180.77-573.15) | 1.64 (0.8) | 2.60±0.35 |

KDT50: 50% Knockdown time in minutes; KDT95: 95% Knockdown time in minutes; CI: Confidence Interval

DDT exhibited prolonged knockdown times, with KDT50 and KDT95 values of 80.14 min and 338.23 min, respectively, in 2021. These values increased slightly in 2022 to 83.17 min and 395.62 min, though they remained lower than the corresponding lambda-cyhalothrin values. Overall, the increases in knockdown times across all insecticides suggest declining knockdown efficacy over time. The probit model provided good fits for all insecticides across both years, with non-significant chi-square values (p > 0.05), except for deltamethrin in 2021 (χ^2^ = 14.18, p = 0.01).

### Resistance Status

The insecticide resistance and susceptibility profiles of *An. gambiae* s.l. populations from UDUS and surrounding villages of Wamako LGA, Sokoto State are presented in **Table 3** and illustrated in **Fig. 5**. In 2021, the population was fully susceptible to alpha-cypermethrin (98%), suspected resistance to deltamethrin (91%), and exhibited resistance to permethrin (89%), lambda-cyhalothrin (76%), and DDT (48%). By 2022, mortality had declined across all insecticides, with resistance confirmed to alpha-cypermethrin (81%), deltamethrin (54%), permethrin (64%), lambda-cyhalothrin (55%), and DDT (41%). These results indicate a clear shift from partial susceptibility to widespread resistance within a year, underscoring the intensifying challenge of managing pyrethroid resistance in local *An. gambiae* s.l. populations.

**Table 3:** Resistance status of *An. gambiae* s.l. populations from Sokoto, north-west Nigeria, exposed to pyrethroids and DDT.

| Year | Insecticide | No. Dead (No. Exposed) | Mortality (%) | Resistance Status |
| --- | --- | --- | --- | --- |
| 2021 | Alpha-cypermethrin (0.05%) | 78 (80) | 98 | Susceptible |
|  | Deltamethrin (0.05%) | 73 (80) | 91 | Possible Resistance |
|  | Permethrin (0.75%) | 71 (80) | 89 | Resistant |
|  | Lambda-cyhalothrin (0.05%) | 76 (100) | 76 | Resistant |
|  | DDT (4%) | 39 (82) | 48 | Resistant |
| 2022 | Alpha-cypermethrin (0.05%) | 65 (80) | 81 | Resistant |
|  | Deltamethrin (0.05%) | 43 (80) | 54 | Resistant |
|  | Permethrin (0.75%) | 51 (80) | 64 | Resistant |
|  | Lambda-cyhalothrin (0.05%) | 44 (80) | 55 | Resistant |
|  | DDT (4%) | 33 (80) | 41 | Resistant |

**Fig. 5:**
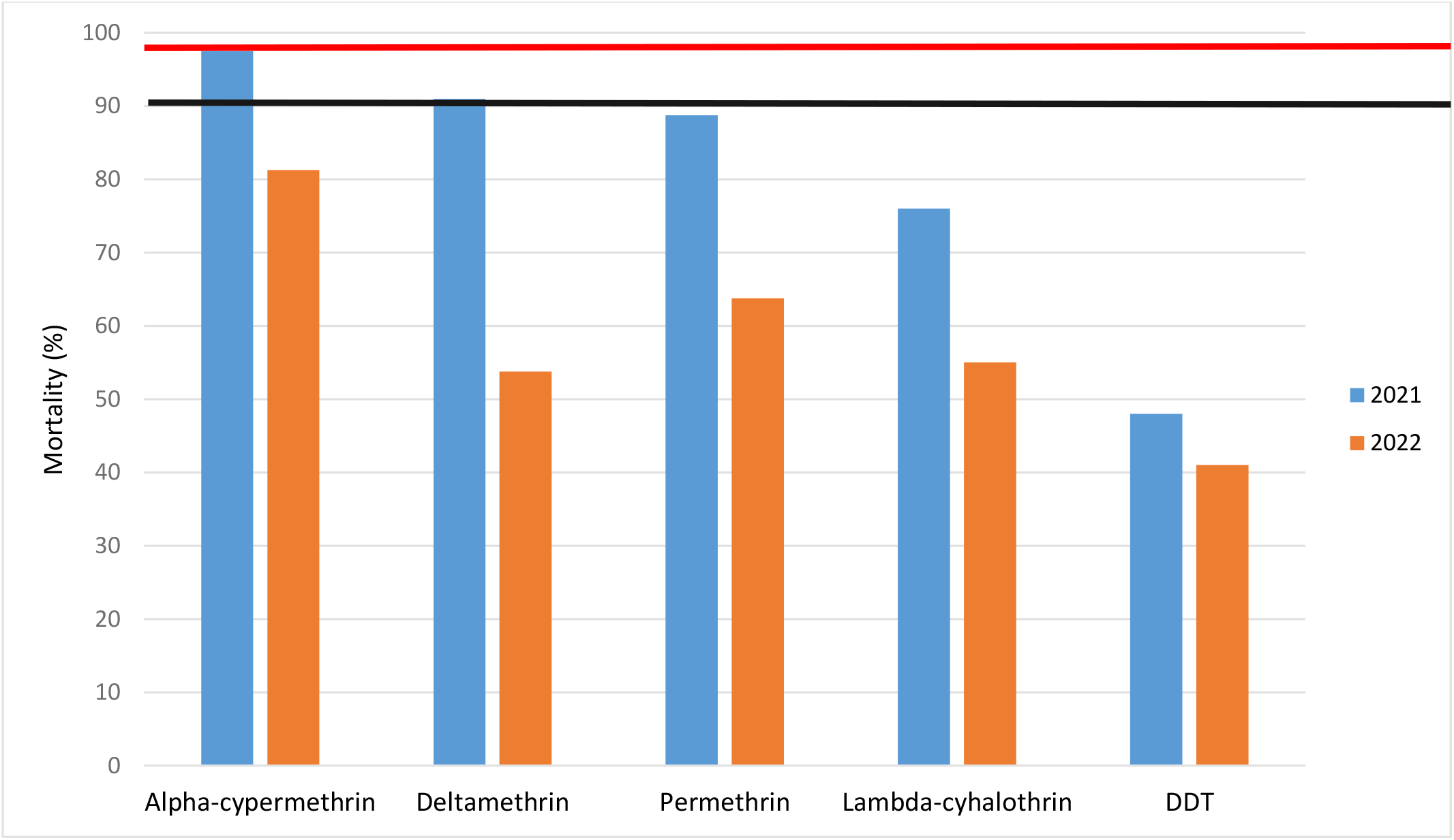
Final (24-hour) Mortality of *An. gambiae* s.l. after exposure to diagnostic concentrations of pyrethroids and DDT using the WHO tube test. The region below the black line represents confirmed resistance; between the black and red lines indicates suspected resistance; and above the red line represents insecticide susceptibility.

### Resistance Intensity

Resistance intensity of *An. gambiae* s.l. to the five insecticides tested using the CDC bottle bioassay at 1×, 5×, and 10× diagnostic doses is presented in **Table 4**. Permethrin and alpha-cypermethrin consistently exhibited low-intensity resistance in both years, with ≥98% mortality at the 5× dose. Lambda-cyhalothrin shifted from low-to moderate-intensity resistance between years, while deltamethrin displayed moderate resistance in 2021 but reduced to low intensity in 2022. In contrast, DDT demonstrated high-intensity resistance in both years, with mortalities of 95% and 90% even at the 10× diagnostic dose.

**Table 4:** Resistance intensity status of *An. gambiae* s.l. after 24-h post exposure to 1x, 5x, and 10x diagnostic concentration of pyrethroids and DDT in the modified CDC bottle bioassay.

| Year | Insecticide | % Mortality (No. Exposed) |  |  | Resistance Intensity Status |
| --- | --- | --- | --- | --- | --- |
|  |  | 1x | 5x | 10x |  |
| 2021 | Alpha-cypermethrin (12.5µg) | 94 (80) | 100 (80) | ND | Low |
|  | DDT (100µg) | 50 (80) | 79 (80) | 95 (80) | High |
|  | Deltamethrin (12.5µg) | 58 (80) | 88 (80) | 100 (80) | Moderate |
|  | Lambda-cyhalothrin (12.5µg) | 74 (80) | 100 (80) | ND | Low |
|  | Permethrin (21.5µg) | 75 (80) | 100 (80) | ND | Low |
| 2022 | Alpha-cypermethrin (12.5µg) | 78 (80) | 98 (80) | ND | Low |
|  | DDT (100µg) | 48 (80) | 73 (80) | 90 (80) | High |
|  | Deltamethrin (12.5µg) | 58 (80) | 100 (80) | ND | Low |
|  | Lambda-cyhalothrin (12.5µg) | 34 (80) | 85 (80) | 100 (80) | Moderate |
|  | Permethrin (21.5µg) | 69 (100) | 100 (100) | ND | Low |

ND, Not done because mortality reached 100% at 5x diagnostic concentration

### Molecular Species Composition of the *An. gambiae* s.l. Complex

The PCR-based identification of *An. gambiae* complex populations revealed the presence of two sibling species: *An. gambiae s*.*s*. and *An. arabiensis* (**Fig. 6**). In 2021, *An. gambiae s*.*s*. accounted for 40% (n=10) of the samples, but its proportion shifted and declined to 18% (n = 4) in 2022. Conversely, *An. arabiensis* constituted the majority in both years, representing 60% (n = 15) in 2021 and increasing its predominance to 82% (n = 18) in 2022.

**Fig. 6:**
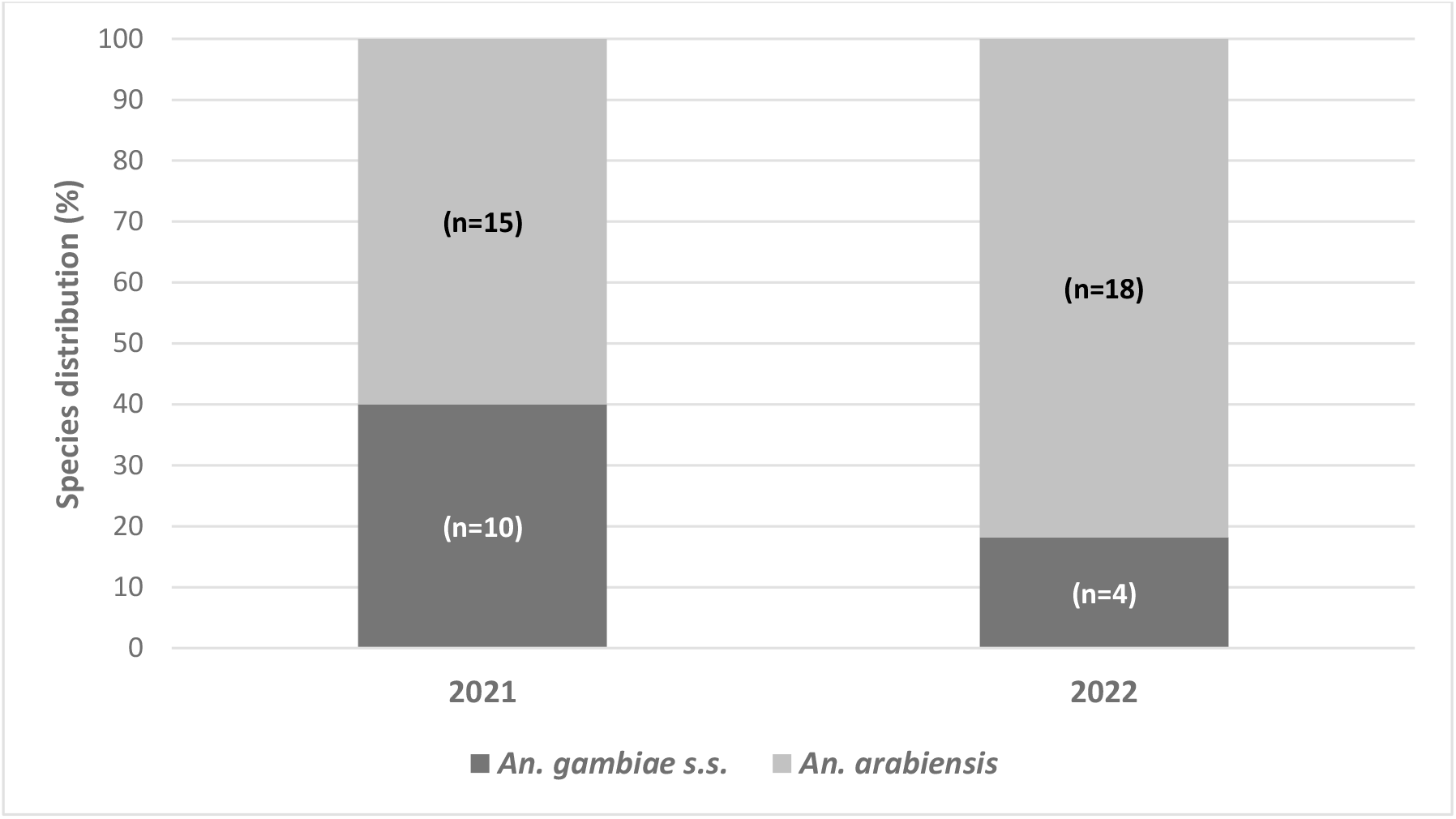
Species composition of the *An. gambiae* s.l. in Sokoto State, Nigeria. The Fig. shows the relative distribution of *An. gambiae s*.*s*. and *An. arabiensis* identified by species-specific PCR in adult An. gambiae s.l. collected in 2021 and 2022. “n” represents the number of individual mosquitoes identified.

### kdr L1014F Mutation Genotyping

Allele-specific PCR targeting the VGSC gene detected the presence of the L1014F mutation among resistant mosquitoes as shown in Table 5. The results demonstrate a consistently high prevalence of kdr L1014F mutation in both sibling species. In 2021, the mean 1014F allele frequency was 0.79, with the highest recorded in *An. gambiae s*.*s*. (0.90), where 80% were F/F. *An. arabiensis* displayed a lower allele frequency (0.67), although most individuals were also F/F = 9. In 2022, *An. gambiae s*.*s*. showed a reduced allele frequency (0.75), with 75% homozygous resistant, while *An. arabiensis* maintained a high frequency (0.75), dominated by the FF genotype (n = 12). H-WE analysis indicated no significant deviation for *An. gambiae s*.*s*. in 2021 (p = 0.7). However, significant deviations from equilibrium were detected in *An. arabiensis* in both years and in *An. gambiae s*.*s*. in 2022 (p < 0.05), suggesting selection pressure acting on the kdr locus.

**Table 5:** Frequencies of kdr L1014F mutations in *An. gambiae* s.s. and *An. arabiensis* populations.

| Year | Species | Number (N) | 1014 kdr genotypes (N) |  |  | 1014F Allele frequency | H-WE *P*-value |
| --- | --- | --- | --- | --- | --- | --- | --- |
|  |  |  | LL | LF | FF |  |  |
| 2021 | *An. gambiae* s.s. | 10 | 0 | 2 | 8 | 0.9 | 0.7 |
|  | *An. arabiensis* | 15 | 4 | 2 | 9 | 0.67 | 0.007 |
| 2022 | *An. gambiae* s.s. | 4 | 1 | 0 | 3 | 0.75 | 0.046 |
|  | *An. arabiensis* | 18 | 3 | 3 | 12 | 0.75 | 0.02 |

LL, homozygous wild type; LF, heterozygous; FF, homozygous mutant.

## Discussion

This study evaluated the insecticide resistance profile of *An. gambiae* s.l. to pyrethroids and DDT over two years (2021–2022), alongside sub-species composition, and the *kdr-w* mutation frequency. The results showed a year-by-year increase in resistance across all insecticides. This trend poses a critical biological threat that undermines the efficacy of LLINs and IRS in Sokoto (49,60). An. gambiae s.l. showed elevated knockdown times to all insecticides. Permethrin and deltamethrin KDT values were substantially higher than those previously reported in Nigeria, indicating reduced susceptibility. Permethrin KDT95 values were 3.4-to 6.8-fold higher, and deltamethrin KDT values were two-to three-fold higher than those reported by (26). DDT produced the highest KDT, consistent with its historic and long-standing resistance (35). These delayed responses serve as early indicators of resistance and likely reflect kdr mutations (61).

Mosquito populations were resistant to DDT, lambda-cyhalothrin, permethrin, and deltamethrin, but initially susceptible to alpha-cypermethrin. However, emerging resistance to alpha-cypermethrin was observed in 2022, consistent with regional trends (62). A similar pattern of moderate-to-high alpha-cypermethrin resistance was also reported in Niger Republic, a region bordering Sokoto State (63). This superior efficacy is attributed to its faster knockdown, strong excito-repellent effects, low cross-resistance, and longer residual activity (64–66). The resistance patterns observed for DDT and pyrethroids are parallel to those documented in both *Anopheles* and *Aedes* species across Nigeria (36,62,67,68). Despite being phased out in the 1980s due to escalating resistance and environmental concerns, DDT continues to show widespread and high resistance, as evident in our study and numerous other reports (30,32,33,37,44,69– 73)

Differences in resistance intensity among pyrethroids is consistent with reports from other high-risk malaria regions (36,74) and pose a serious threat to malaria control in Nigeria (75). This variation among insecticides of same class further suggests contributions from metabolic resistance mechanisms, supported by observed shifts in resistance profiles and the likely involvement of detoxification enzymes or kdr allele selection (29,76,77).

Molecular identification detected two sibling species, *An. gambiae s*.*s*. and *An. arabiensis*, with the latter predominating. Its dominance aligns with established patterns in arid and semi-arid regions of Nigeria (78–81). In contrast, some authors documented the predominance of *An. coluzzii* in similar Sudano-Sahelian transects (30,82), although no *An. coluzzii* were detected in our study, likely due to methodological limitations and sample size constraints. This observed shift towards *An. arabiensis* dominance is consistent with recent findings across West and East Africa, particularly in urban and peri-urban environments (83,84). The widespread deployment of indoor vector control interventions may selectively disadvantage endophilic species, indirectly favouring exophilic and exophagic vectors such as An. arabiensis (85). Furthermore, larval sampling, which favours detection of this species, and extensive livestock rearing in surrounding communities may have reinforced its predominance, as *An. arabiensis* is highly zoophilic (86–88).

A high frequency of the L1014F mutation was detected, particularly in *An. gambiae s*.*s*., consistent with reports from Nigeria and Niger republic (29,63). This mutation likely accumulates over time with continued insecticide use. For example, the previously reported low frequency of 13.5% in northern Nigeria in 2014 (29) contrasts with the substantially higher prevalence observed in our study population. Nevertheless, our values remain lower than the highest frequency reported from both low- and high-risk villages in Ethiopia (89). Beyond resistance, *kdr* mutation has also been associated with increase mosquito susceptibility to *Plasmodium* infection, thereby potentially amplifying malaria transmission (90,91). The observed deviation from H–WE suggest strong selection pressure at the kdr locus, favoring resistant homozygotes under continued pyrethroid pressure (61,89). Factors such as regional movement and trade may further facilitate the spread of kdr-resistant mosquitoes, posing challenges not only for Nigeria but also for neighboring regions (92,93). The limitations of our study include the restricted geographic scope and sample size, the absence of synergist bioassays e.g., piperonyl butoxide (PBO) to assess the role of metabolic resistance mechanisms, and the lack of multi-locus genotyping to provide a more comprehensive understanding of resistance dynamics.

## Conclusions

In conclusion, this study underscores the critical threat of insecticide resistance in *An. gambiae* populations in Sokoto, Northwestern Nigeria, characterized by high resistance to DDT and low-to-high resistance to permethrin, deltamethrin, alpha-cypermethrin, and lambda-cyhalothrin. The elevated knockdown times and high prevalence of the kdr-w L1014F mutation highlight a concerning escalation of resistance with a strong genetic basis. These findings emphasize the urgent need for integrated vector and targeted resistance management strategies, including rotation, combination strategies, or alternative insecticides to sustain malaria control efforts in Northern Nigeria.

## Abbreviations

AA: Amino acid
AS-PCR: Allele-specific polymerase chain reaction
CDC: Centers for Disease Control and Prevention
DDT: Dichlorodiphenyltrichloroethane
F/F: Homozygous resistant
GEIS: Global Emerging Infections Surveillance
H–WE: Hardy–Weinberg equilibrium
IGS: Intergenic spacer
IRS: Indoor residual spraying
ITNs: Insecticide-treated nets
KDT: Knockdown time
KDT50: Time to 50% knockdown
KDT95: Time to 95% knockdown
kdr: Knockdown resistance
kdr-w: West African knockdown resistance mutation (L1014F)
L/L: Homozygous susceptible
L/F: Heterozygous
LLINs: Long-lasting insecticidal nets
NEMC: Nigeria End Malaria Council
NFW: Nuclease-free water
NMEP: National Malaria Elimination Programme
NMSP: National Malaria Strategic Plan
PBO: Piperonyl butoxide
PCR: Polymerase chain reaction
PMI: President’s Malaria Initiative
s.l.: *Sensu lato*
s.s.: *Sensu stricto*
UDUS: Usmanu Danfodiyo University Sokoto
VGSC: Voltage-gated sodium channel
WHO: World Health Organization
WRBU: Walter Reed Biosystematics Unit

## Declarations

### Ethics approval and consent to participate

Not applicable

### Consent for publication

Not applicable.

### Availability of data and materials

All data generated or analysed during this study are included in this published article.

## Acknowledgements

We extend our sincere gratitude to the inhabitants of the communities at the various collection sites and the VectorLink USPMI Sokoto sentinel team for their support during the sample collection and insecticide bioassays.

## Disclaimer

Material has been reviewed by the authors’ respective institutions. There is no objection to its presentation and/or publication. The opinions or assertions contained herein are the private views of the author, and are not to be construed as official, or as reflecting true views of the Department of the Army or the Department of Defense.

## References

1. World Health Organization. World Malaria Report 2023 [Internet]. World Health Organization; 2023 [cited 2025 Sep 29]. Available from: Licence: CC BY-NC-SA 3.0 IGO.

2. Bhuvaneswari A, Shriram AN, Raju KHK, Kumar A. Mosquitoes, Lymphatic Filariasis, and Public Health: A Systematic Review of Anopheles and Aedes Surveillance Strategies. Pathogens. 2023 Nov 29;12(12):1406.

3. Okorie PN, McKenzie FE, Ademowo OG, Bockarie M, Kelly-Hope L. Nigeria Anopheles Vector Database: An Overview of 100 Years’ Research. PLoS One. 2011 Dec 5;6(12):e28347.

4. Adeogun AO, Babalola AS, Oyale OO, Oyeniyi T, Omotayo A, Izekor RT, et al. Spatial distribution and geospatial modeling of potential spread of secondary malaria vectors species in Nigeria using recently collected empirical data. PLoS One. 2025 Apr 21;20(4):e0320531.

5. World Health Organization. World malaria report 2024: addressing inequity in the global malaria response. Geneva: World Health Organization [Internet]. World Health Organization; 2024 [cited 2025 Sep 29]. Available from: Licence: CC BY-NC-SA 3.0 IGO.

6. National Malaria Elimination Programme FM of HN. NATIONAL MALARIA STRATEGIC PLAN, 2021 – 2025 [Internet]. 2020. Available from: https://nmcp.gov.ng

7. World Health Organization. Report on malaria in Nigeria 2022. World Health Organization, Regional Office for Africa; 2023. 87 p.

8. U.S. President’s Malaria Initiative. U.S. President’s Malaria Initiative Nigeria Malaria Operational Plan FY 2022 [Internet]. 2022. Available from: www.pmi.gov

9. Omojuyigbe JO, Owolade AJJ, Sokunbi TO, Bakenne HA, Ogungbe BA, Oladipo HJ, et al. Malaria eradication in Nigeria: State of the nation and priorities for action. Journal of Medicine, Surgery, and Public Health. 2023;1:100024.

10. World Health Organization. https://www.who.int/news/item/15-08-2007-who-releases-new-guidance-on-insecticide-treated-mosquito-nets. 2007. WHO releases new guidance on insecticide-treated mosquito nets.

11. Benelli G, Beier JC. Current vector control challenges in the fight against malaria. Acta Trop. 2017 Oct;174:91–6.

12. van den Berg H, da Silva Bezerra HS, Al-Eryani S, Chanda E, Nagpal BN, Knox TB, et al. Recent trends in global insecticide use for disease vector control and potential implications for resistance management. Sci Rep. 2021 Dec 13;11(1):23867.

13. Kleinschmidt I, Bradley J, Knox TB, Mnzava AP, Kafy HT, Mbogo C, et al. Implications of insecticide resistance for malaria vector control with long-lasting insecticidal nets: a WHO-coordinated, prospective, international, observational cohort study. Lancet Infect Dis. 2018 Jun;18(6):640–9.

14. Gueye A, Ngom EHM, Ndoye BB, Dione ML, Diouf B, Ndiaye EH, et al. Insecticide Resistance and Target-Site Mutations kdr, N1575Y, and Ace-1 in Anopheles gambiae s.l. Populations in a Low-Malaria-Transmission Zone in the Sudanian Region of Senegal. Genes (Basel). 2024 Oct 16;15(10):1331.

15. Oyeniyi AT, Adeogun AO, Idowu ET, Oboh B, Olakiigbe A, Adesalu O, et al. First report of N1575Y mutation in pyrethroid resistant Anopheles gambiae s.l. in Nigeria. Sci Afr. 2020 Nov;10:e00645.

16. Fanello C, Petrarca V, Della Torre A, Santolamazza F, Dolo G, Coulibaly M, et al. The pyrethroid knock-down resistance gene in the Anopheles gambiae complex in Mali and further indication of incipient speciation within An. gambiae s.s. Insect Mol Biol. 2003 Jun 9;12(3):241–5.

17. Yawson AE, Mccall PJ, Wilson MD, Donnelly MJ. Species abundance and insecticide resistance of Anopheles gambiae in selected areas of Ghana and Burkina Faso. Med Vet Entomol. 2004 Dec 7;18(4):372–7.

18. Keïta M, Sogoba N, Kané F, Traoré B, Zeukeng F, Coulibaly B, et al. Multiple Resistance Mechanisms to Pyrethroids Insecticides in Anopheles gambiae sensu lato Population From Mali, West Africa. J Infect Dis. 2021 Apr 27;223(Supplement_2):S81–90.

19. Zuharah WF, Sufian M. The discovery of a novel knockdown resistance (kdr) mutation A1007G on Aedes aegypti (Diptera: Culicidae) from Malaysia. Sci Rep. 2021 Mar 4;11(1):5180.

20. Singh OP, Dykes CL, Das MK, Pradhan S, Bhatt RM, Agrawal OP, et al. Presence of two alternative kdr-like mutations, L1014F and L1014S, and a novel mutation, V1010L, in the voltage gated Na+ channel of Anopheles culicifacies from Orissa, India. Malar J. 2010 Dec 28;9(1):146.

21. Silva APB, Santos JMM, Martins AJ. Mutations in the voltage-gated sodium channel gene of anophelines and their association with resistance to pyrethroids – a review. Parasit Vectors. 2014 Dec 7;7(1):450.

22. Bass C, Nikou D, Donnelly MJ, Williamson MS, Ranson H, Ball A, et al. Detection of knockdown resistance (kdr) mutations in Anopheles gambiae: a comparison of two new high-throughput assays with existing methods. Malar J. 2007 Dec 13;6(1):111.

23. Bruce-Chwatt LJ, Haworth J. Annual Report of the Malarial Control Pilot Project in Western Sokoto (Northern Nigeria) 1955–56. International Journal of Pest Management: Part A. 1957 Feb;3(1):18–22.

24. Service MW. The behaviour of malaria vectors in huts sprayed with DDT and with a mixture of DDT and malathion in Northern Nigeria. Trans R Soc Trop Med Hyg. 1964 Jan;58(1):72–9.

25. Awolola TS, Brooke BD, Hunt RH, Coetze M. Resistance of the malaria vector Anopheles gambiae s.s. to pyrethroid insecticides, in south-western Nigeria. Ann Trop Med Parasitol. 2002 Dec 1;96(8):849–52.

26. Awolola TS, Adeogun A, Olakiigbe AK, Oyeniyi T, Olukosi YA, Okoh H, et al. Pyrethroids resistance intensity and resistance mechanisms in Anopheles gambiae from malaria vector surveillance sites in Nigeria. PLoS One. 2018 Dec 5;13(12):e0205230.

27. Kayode IF, Emmanuel Taiwo I, Adedapo O A, Olalekan O, Precious Chimdalu I, Iyanuoluwa Olayilola O, et al. Low frequency of knockdown resistance mutation (L1014F) and the efficacy of PBO synergist in multiple insecticide-resistant populations of Anopheles gambiae in Ikorodu, Lagos State, Nigeria. Afr Health Sci. 2023 Apr 6;23(1):255–61.

28. Djouaka R, Riveron JM, Yessoufou A, Tchigossou G, Akoton R, Irving H, et al. Multiple insecticide resistance in an infected population of the malaria vector Anopheles funestus in Benin. Parasit Vectors. 2016 Dec 17;9(1):453.

29. Ibrahim SS, Manu YA, Tukur Z, Irving H, Wondji CS. High frequency of kdr L1014F is associated with pyrethroid resistance in Anopheles coluzzii in Sudan savannah of northern Nigeria. BMC Infect Dis. 2014 Dec 15;14(1):441.

30. Ibrahim SS, Mukhtar MM, Datti JA, Irving H, Kusimo MO, Tchapga W, et al. Temporal escalation of Pyrethroid Resistance in the major malaria vector Anopheles coluzzii from Sahelo-Sudanian Region of northern Nigeria. Sci Rep. 2019 May 14;9(1):7395.

31. Ibrahim SS, Mukhtar MM, Irving H, Riveron JM, Fadel AN, Tchapga W, et al. Exploring the Mechanisms of Multiple Insecticide Resistance in a Highly Plasmodium-Infected Malaria Vector Anopheles funestus Sensu Stricto from Sahel of Northern Nigeria. Genes (Basel). 2020 Apr 22;11(4):454.

32. Wahedi JA, Ande AT, Oduola AO, Obembe A. Bendiocarb resistance and kdr associated deltamethrin and DDT resistance in Anopheles gambiae s.l. populations from North Eastern Adamawa State, Nigeria. Ceylon Journal of Science. 2021 Mar 15;50(1):63.

33. Olatunbosun-Oduola A, Abba E, Adelaja O, Taiwo-Ande A, Poloma-Yoriyo K, Samson-Awolola T. Widespread Report of Multiple Insecticide Resistance in Anopheles gambiae s.l. Mosquitoes in Eight Communities in Southern Gombe, North-Eastern Nigeria. J Arthropod Borne Dis. 2019 Mar;13(1):50–61.

34. Kristan M, Fleischmann H, Della Torre A, Stich A, Curtis CF. Pyrethroid resistance/susceptibility and differential urban/rural distribution of Anopheles arabiensis and An. gambiae s.s. malaria vectors in Nigeria and Ghana. Med Vet Entomol. 2003 Sep 27;17(3):326–32.

35. Fagbohun IK, Oyeniyi TA, Idowu TE, Otubanjo OA, Awolola ST. Cytochrome P450 Mono-Oxygenase and Resistance Phenotype in DDT and Deltamethrin-Resistant Anopheles gambiae (Diptera: Culicidae) and Culex quinquefasciatus in Kosofe, Lagos, Nigeria. J Med Entomol. 2019 Apr 16;56(3):817–21.

36. Muhammad A, Ibrahim SS, Mukhtar MM, Irving H, Abajue MC, Edith NMA, et al. High pyrethroid/DDT resistance in major malaria vector Anopheles coluzzii from Niger-Delta of Nigeria is probably driven by metabolic resistance mechanisms. PLoS One. 2021 Mar 11;16(3):e0247944.

37. Habibu UA, Yayo AM, Yusuf YD. Susceptibility Status Of Anopheles Gambiae Complex To Insecticides Commonly Used For Malaria Control In Northern Nigeria. INTERNATIONAL JOURNAL OF SCIENTIFIC & TECHNOLOGY RESEARCH. 2017;6(06).

38. Oduola AO, Idowu ET, Oyebola MK, Adeogun AO, Olojede JB, Otubanjo OA, et al. Evidence of carbamate resistance in urban populations of Anopheles gambiae s.s. mosquitoes resistant to DDT and deltamethrin insecticides in Lagos, South-Western Nigeria. Parasit Vectors. 2012 Dec 11;5(1):116.

39. Okorie PN, Ademowo OG, Irving H, Kelly-HOPE LA, Wondji CS. Insecticide susceptibility of Anopheles coluzzii and Anopheles gambiae mosquitoes in Ibadan, Southwest Nigeria. Med Vet Entomol. 2015 Mar 22;29(1):44–50.

40. Adeogun AO, Popoola KO, Oduola AO, Olakiigbe AK, Awolola ST. High Level of DDT Resistance and Reduced Susceptibility to Deltamethrin in Anopheles gambiae, Anopheles coluzzi, and Anopheles arabiensis from Urban Communities in Oyo State, South-West Nigeria. J Mosq Res. 2017;

41. Santolamazza F, Calzetta M, Etang J, Barrese E, Dia I, Caccone A, et al. Distribution of knock-down resistance mutations in Anopheles gambiae molecular forms in west and west-central Africa. Malar J. 2008 Dec 29;7(1):74.

42. Omotayo AI, Dogara MM, Sufi D, Shuaibu T, Balogun J, Dawaki S, et al. High pyrethroid-resistance intensity in Culex quinquefasciatus (Say) (Diptera: Culicidae) populations from Jigawa, North-West, Nigeria. PLoS Negl Trop Dis. 2022 Jun 21;16(6):e0010525.

43. Ekedo CM, Ukpai OM, Ehisianya CN, Nwangwu UC, Nwosu EM, Adeogun AO, et al. Insecticide resistance spectrum and prevalence of L1014F kdr type mutation in Anopheles gambiae s.l. in Abia State, Nigeria. Ceylon Journal of Science. 2023 Jun 1;52(2):163–74.

44. Chukwuekezie O, Nwosu E, Nwangwu U, Dogunro F, Onwude C, Agashi N, et al. Resistance status of Anopheles gambiae (s.l.) to four commonly used insecticides for malaria vector control in South-East Nigeria. Parasit Vectors. 2020 Dec 24;13(1):152.

45. Safiyanu M Aayaisia and AH. Detection of KDR l1014f mutation in pyrethroids susceptible Anopheles gambiae S.L from Ladanai, Kano state, northwest Nigeria. Int J Mosq Res. 2019;6(3):10–5.

46. PMI VectorLink Project. PMI VectorLink Project Annual Report October 1, 2020 – September 30, 2021 [Internet]. 2021 Nov. Available from: www.abtassociates.com

47. Obembe A, Oduola AO, Adeogun A, Inyang U, Oyeniyi T, Olakiigbe A, et al. Implementation of malaria vector surveillance and insecticide resistance monitoring interventions in Nigeria. Glob Health Res Policy. 2024 Dec 31;9(1):55.

48. Implications of Insecticide Resistance Consortium. Implications of insecticide resistance for malaria vector control with long-lasting insecticidal nets: trends in pyrethroid resistance during a WHO-coordinated multi-country prospective study. Parasit Vectors. 2018 Dec 22;11(1):550.

49. World Health Organization. Global report on insecticide resistance in malaria vectors: 2010–2016 [Internet]. 2018 [cited 2025 Sep 30]. Available from: https://www.who.int/publications/i/item/9789241514057

50. Service MW. Mosquito Ecology : Field Sampling Methods. 2001;998.

51. Parker C. Collection and Rearing of Container Mosquitoes and a 24-h Addition to the CDC Bottle Bioassay. Journal of Insect Science [Internet]. 2020 Nov 1 [cited 2025 Nov 9];20(6):13. Available from: https://pmc.ncbi.nlm.nih.gov/articles/PMC7751146/

52. Abdullahi YM, Fana SA, Umar YS, Batagarawa US. Prevalence of Mosquitoes in Gidan Yunfa Community of Usmanu Danfodiyo University, Sokoto, Nigeria. Path of Science [Internet]. 2020 Jan 1 [cited 2025 Nov 9];6(5):8001–6. Available from: https://www.academia.edu/63650913/Prevalence_of_Mosquitoes_in_Gidan_Yunfa_Community_of_Usmanu_Danfodiyo_University_Sokoto_Nigeria

53. Tsegaye A, Demissew A, Hawaria D, Abossie A, Getachew H, Habtamu K, et al. Anopheles larval habitats seasonality and environmental factors affecting larval abundance and distribution in Arjo-Didessa sugar cane plantation, Ethiopia. Malar J [Internet]. 2023 Dec 1 [cited 2025 Nov 9];22(1):350. Available from: https://pmc.ncbi.nlm.nih.gov/articles/PMC10652594/

54. World Health Organization. Test procedures for insecticide resistance monitoring in malaria vector mosquitoes (Second edition) (Updated June 2018). Who [Internet]. 2018 [cited 2025 Sep 30];48. Available from: http://www.who.int/malaria/publications/atoz/9789241511575/en/

55. Gillies MT., Coetzee Maureen. A supplement to the anophelinae of Africa south of the Sahara (Afrotropical Region). South African Institute for Medical Research; 1987. 143 p.

56. CDC. Guideline for Evaluating Insecticide Resistance in Vectors Using the CDC Bottle Bioassay [Internet]. 2012. Available from: http://www.cdc.gov/malaria.

57. Scott JA, Brogdon WG, Collins FH. Identification of single specimens of the Anopheles gambiae complex by the polymerase chain reaction. Am J Trop Med Hyg [Internet]. 1993 [cited 2025 Sep 30];49(4):520–9. Available from: https://pubmed.ncbi.nlm.nih.gov/8214283/

58. Martinez-Torres D, Chandre F, Williamson MS, Darriet F, Bergé JB, Devonshire AL, et al. Molecular characterization of pyrethroid knockdown resistance (kdr) in the major malaria vector Anopheles gambiae s.s. Insect Mol Biol [Internet]. 1998 May [cited 2025 Sep 30];7(2):179–84. Available from: https://pubmed.ncbi.nlm.nih.gov/9535162/

59. Finney DJ. Probit Analysis. J Pharm Sci [Internet]. 1971 Sep 1 [cited 2025 Sep 30];60(9):1432. Available from:/doi/pdf/10.1002/jps.2600600940

60. Chandre F, Darriet F, Duchon S, Finot L, Manguin S, Carnevale P, et al. Modifications of pyrethroid effects associated with kdr mutation in Anopheles gambiae. Med Vet Entomol. 2000 Mar 25;14(1):81–8.

61. Abdullahi YM, Fana SA, Bandiya HM, Batagarawa US, Zulkarnain M, Uthman HL. Susceptibility status of Aedes aegypti to pyrethroids in Usmanu Danfodiyo University main campus, Sokoto, Nigeria. Int J Mosq Res. 2022 Mar 1;9(2):30–6.

62. Soumaila H, Hamani B, Arzika II, Soumana A, Daouda A, Daouda FA, et al. Countrywide insecticide resistance monitoring and first report of the presence of the L1014S knock down resistance in Niger, West Africa. Malar J. 2022 Dec 16;21(1):385.

63. Abbas N, Hafez AM. Alpha-Cypermethrin Resistance in Musca domestica: Resistance Instability, Realized Heritability, Risk Assessment, and Insecticide Cross-Resistance. Insects. 2023 Feb 26;14(3):233.

64. Uragayala S, Kamaraju R, Tiwari S, Ghosh SK, Valecha N. Small-scale evaluation of the efficacy and residual activity of alpha-cypermethrin WG (250 g AI/kg) for indoor spraying in comparison with alpha-cypermethrin WP (50 g AI/kg) in India. Malar J. 2015 Dec 29;14(1):223.

65. Ngufor C, Agbevo A, Fagbohoun J, Fongnikin A, Rowland M. Efficacy of Royal Guard, a new alpha-cypermethrin and pyriproxyfen treated mosquito net, against pyrethroid-resistant malaria vectors. Sci Rep. 2020 Jul 22;10(1):12227.

66. Adamu AS, Suleiman M, Sani I. Identification and assessment of knock down rate of Aedes mosquitoes in northern Katsina state, Nigeria. Int J Mosq Res. 2023 Jan 1;10(6):111–6.

67. Abdullahi YM, Fana SA, Batagarawa US. Efficacy of Chlorfenapyr and Clothianidin Insecticides against Permethrin Resistant Anopheles gambiae s.l. in Gidan Yaro Village, Sokoto, Nigeria. Asian Journal of Biological Sciences. 2022 Jul 1;15(4):227–34.

68. Nutifafa GG, Hanafi-Bojd AA, Oshaghi M, Dadzie S, Vatandoost H, Koosha M. Insecticide Susceptibility status of An. gambiae s.l. (Culicidae: Giles) from selected in-land and coastal agricultural areas of Ghana. J Entomol Zool Stud. 2017;5(1):701–7.

69. Boussougou-Sambe ST, Ngossanga B, Doumba-Ndalembouly AG, Boussougou LN, Woldearegai TG, Mougeni F, et al. Anopheles gambiae s.s. resistance to pyrethroids and DDT in semi-urban and rural areas of the Moyen-Ogooué Province Gabon. Malar J. 2023 Dec 1;22(1).

70. Dhiman S, Yadav K, Rabha B, Goswami D, Hazarika S, Tyagi V. Evaluation of Insecticides Susceptibility and Malaria Vector Potential of Anopheles annularis s.l. and Anopheles vagus in Assam, India. PLoS One. 2016 Mar 24;11(3):e0151786.

71. Abbasi E, Vahedi M, Bagheri M, Gholizadeh S, Alipour H, Moemenbellah-Fard MD. Monitoring of synthetic insecticides resistance and mechanisms among malaria vector mosquitoes in Iran: A systematic review. Heliyon. 2022 Jan;8(1):e08830.

72. Orjuela LI, Morales JA, Ahumada ML, Rios JF, González JJ, Yañez J, et al. Insecticide Resistance and Its Intensity in Populations of Malaria Vectors in Colombia. Biomed Res Int. 2018 Aug 29;2018:1–12.

73. Pwalia R, Joannides J, Iddrisu A, Addae C, Acquah-Baidoo D, Obuobi D, et al. High insecticide resistance intensity of Anopheles gambiae (s.l.) and low efficacy of pyrethroid LLINs in Accra, Ghana. Parasit Vectors. 2019 Dec 13;12(1):299.

74. Yusuf MA, Oshaghi MA, Vatandoost H, Hanafi-Bojd AA, Enayati A, Jalo RI, et al. Current Status of Insecticide Susceptibility in the Principal Malaria Vector, Anopheles gambiae in Three Northern States of Nigeria. J Arthropod Borne Dis. 2021 Oct 17;

75. Djouaka RF, Bakare AA, Coulibaly ON, Akogbeto MC, Ranson H, Hemingway J, et al. Expression of the cytochrome P450s, CYP6P3 and CYP6M2 are significantly elevated in multiple pyrethroid resistant populations of Anopheles gambiae s.s. from Southern Benin and Nigeria. BMC Genomics. 2008 Dec 13;9(1):538.

76. Witzig C, Parry M, Morgan JC, Irving H, Steven A, Cuamba N, et al. Genetic mapping identifies a major locus spanning P450 clusters associated with pyrethroid resistance in kdr-free Anopheles arabiensis from Chad. Heredity (Edinb). 2013 Apr 9;110(4):389–97.

77. Onyabe DY, Conn JE. The distribution of two major malaria vectors, Anopheles gambiae and Anopheles arabiensis, in Nigeria. Mem Inst Oswaldo Cruz. 2001 Nov;96(8):1081–4.

78. Sinka ME, Bangs MJ, Manguin S, Coetzee M, Mbogo CM, Hemingway J, et al. The dominant Anopheles vectors of human malaria in Africa, Europe and the Middle East: occurrence data, distribution maps and bionomic précis. Parasit Vectors. 2010 Dec 3;3(1):117.

79. Lindsay SW, Parson L, Thomas CJ. Mapping the range and relative abundance of the two principal African malaria vectors, Anopheles gambiae sensu stricto and An. arabiensis, using climate data. Proc R Soc Lond B Biol Sci. 1998 May 22;265(1399):847–54.

80. Coetzee M, Craig M, le Sueur D. Distribution of African Malaria Mosquitoes Belonging to the Anopheles gambiae Complex. Parasitology Today. 2000 Feb;16(2):74–7.

81. Ibrahim SS, Muhammad A, Hearn J, Weedall GD, Nagi SC, Mukhtar MM, et al. Molecular drivers of insecticide resistance in the Sahelo-Sudanian populations of a major malaria vector Anopheles coluzzii. BMC Biol. 2023 May 24;21(1):125.

82. Iga J, Ochaya S, Echodu R, Opiyo EA, Musiime AK, Nakamaanya A, et al. Sibling Species Composition and Susceptibility Status of Anopheles gambiae s.l. to Insecticides Used for Indoor Residual Spraying in Eastern Uganda. J Parasitol Res. 2023 Jul 10;2023:1–8.

83. Mwesige R, Byagamy JP, Opiro R, Angwech H, Onanyang D, Ocen PB, et al. Sibling species composition and feeding pattern of malaria vectors in indoor-sprayed and non-sprayed districts of Lira and Kole, northern Uganda. Malar J. 2025 Jul 1;24(1):202.

84. Abeku TA, Helinski MEH, Kirby MJ, Ssekitooleko J, Bass C, Kyomuhangi I, et al. Insecticide resistance patterns in Uganda and the effect of indoor residual spraying with bendiocarb on kdr L1014S frequencies in Anopheles gambiae s.s. Malar J. 2017 Apr 20;16(1):156.

85. Ebhodaghe FI, Sanchez-Vargas I, Isaac C, Foy BD, Hemming-Schroeder E. Sibling species of the major malaria vector Anopheles gambiae display divergent preferences for aquatic breeding sites in southern Nigeria. Malar J. 2024 Feb 27;23(1):60.

86. Asale A, Duchateau L, Devleesschauwer B, Huisman G, Yewhalaw D. Zooprophylaxis as a control strategy for malaria caused by the vector Anopheles arabiensis (Diptera: Culicidae): a systematic review. Infect Dis Poverty. 2017 Oct 25;6(1):160.

87. Gone T, Balkew M, Gebre-Michael T. Comparative entomological study on ecology and behaviour of Anopheles mosquitoes in highland and lowland localities of Derashe District, southern Ethiopia. Parasit Vectors. 2014 Dec 20;7(1):483.

88. Yewhalaw D, Bortel W Van, Denis L, Coosemans M, Duchateau L, Speybroeck N. First Evidence of High Knockdown Resistance Frequency in Anopheles arabiensis (Diptera: Culicidae) from Ethiopia. The American Society of Tropical Medicine and Hygiene. 2010 Jul;83(1):122–5.

89. Alout H, Ndam NT, Sandeu MM, Djégbe I, Chandre F, Dabiré RK, et al. Insecticide Resistance Alleles Affect Vector Competence of Anopheles gambiae s.s. for Plasmodium falciparum Field Isolates. PLoS One. 2013 May 21;8(5):e63849.

90. Alout H, Dabiré RK, Djogbénou LS, Abate L, Corbel V, Chandre F, et al. Interactive cost of Plasmodium infection and insecticide resistance in the malaria vector Anopheles gambiae. Sci Rep. 2016 Jul 19;6(1):29755.

91. Aïzoun N, Aïkpon R, Akogbéto M. Evidence of increasing L1014F kdr mutation frequency in Anopheles gambiae s.l. pyrethroid resistant following a nationwide distribution of LLINs by the Beninese National Malaria Control Programme. Asian Pac J Trop Biomed. 2014 Mar;4(3):239–43.

92. World Health Organization. Effectiveness of disinsection of conveyances to prevent or reduce the spread of mosquito vectors via international travel. World Health Organization; 2024.

